# Multi-Trait Improvement by Predicting Genetic Correlations in Breeding Crosses

**DOI:** 10.1101/593210

**Authors:** Jeffrey L. Neyhart, Aaron J. Lorenz, Kevin P. Smith

**Affiliations:** Department of Agronomy and Plant Genetics, University of Minnesota, St. Paul, MN 55108

**Keywords:** genetic correlation, genomewide prediction, cross selection, simulation, barley

## Abstract

The many quantitative traits of interest to plant breeders are often genetically correlated, which can complicate progress from selection. Improving multiple traits may be enhanced by identifying parent combinations – an important breeding step – that will deliver more favorable genetic correlations (*r*_*G*_). Modeling the segregation of genomewide markers with estimated effects may be one method of predicting *r*_*G*_ in a cross, but this approach remains untested. Our objectives were to: (i) use simulations to assess the accuracy of genomewide predictions of *r*_*G*_ and the long-term response to selection when selecting crosses on the basis of such predictions; and (ii) empirically measure the ability to predict genetic correlations using data from a barley (*Hordeum vulgare* L.) breeding program. Using simulations, we found that the accuracy to predict *r*_*G*_ was generally moderate and influenced by trait heritability, population size, and genetic correlation architecture (i.e. pleiotropy or linkage disequilibrium). Among 26 barley breeding populations, the empirical prediction accuracy of *r*_*G*_ was low (−0.012) to moderate (0.42), depending on trait complexity. Within a simulated plant breeding program employing indirect selection, choosing crosses based on predicted *r*_*G*_ increased multi-trait genetic gain by 11-27% compared to selection on the predicted cross mean. Importantly, when the starting genetic correlation was negative, such cross selection mitigated or prevented an unfavorable response in the trait under indirect selection. Prioritizing crosses based on predicted genetic correlation can be a feasible and effective method of improving unfavorably correlated traits in breeding programs.

## INTRODUCTION

Quantitative traits often exhibit complex relationships with one another, with ramifications for disease epidemiology, evolutionary processes, and plant and animal improvement. These relationships may manifest as genetic correlations, which can be caused by shared genetic influence (i.e. pleiotropy) or the non-random association of alleles (i.e. linkage disequilibrium) (Lynch and Walsh 1998). Investigations in quantitative genetics commonly assume that many loci of small effect govern traits [i.e. “infinitesimal model” (Fisher 1919)]. This suggest that a large proportion of the genome should contribute to phenotypic variation, a hypothesis that has been supported by recent genome-wide analyses of complex traits (Mackay 2010; Boyle *et al.* 2017). If true for multiple complex traits, a natural corollary follows that pleiotropy or close linkage of trait-specific genes is widespread. Recent studies attempting to identify quantitative trait loci (QTL) influencing multiple traits using dense genomewide markers have provided support for this idea, reporting extensive pleiotropy or strong genetic correlations (Korte *et al.* 2012; Lee *et al.* 2012; Bulik-Sullivan *et al.* 2015; Schaid *et al.* 2016; Deng and Pan 2017).

Plant breeders routinely select on multiple traits, but progress can be complicated by genetic correlations. If two traits are favorably correlated, selection can simultaneously improve both by tandem selection, indirect selection, or a trait index (Bernardo 2010). Unfavorable correlations, meanwhile, are common and often the bane of the breeder. In crop improvement, notorious examples include grain yield and grain protein content in wheat (*Triticum aestivum* L.; Simmonds 1995), grain yield and plant height in maize (*Zea mays* L.; Chi *et al.* 1969), and seed protein and oil content in soybean (*Glycine max* L.; Bandillo *et al.* 2015). The directions of such correlations imply an unfavorable response in one trait when selecting on another (Falconer and Mackay 1996), and the underlying cause will impact the prospects of long-term improvement. Selection on traits with shared, antagonistic genetic influence is functionally constrained, but correlations induced by linkage disequilibrium are transient and can be disrupted by recombination (Falconer and Mackay 1996; Lynch and Walsh 1998).

Genomewide selection has become popular among plant breeders as a method of predicting the merit of unphenotyped individuals using genomewide markers and a phenotyped training population (Meuwissen *et al.* 2001). Typical prediction models are univariate (i.e. one trait), but multivariate models have recently been explored as a means of borrowing information from genetically correlated traits and improving the prediction accuracy of both traits (Calus and Veerkamp 2011; Jia and Jannink 2012). Selection on multiple traits using predicted breeding values would proceed as if using phenotypic values, relying on procedures such as tandem selection, independent culling levels, or the construction of a trait index (Bernardo 2010), with most studies of multi-trait genomewide selection using the latter (Combs and Bernardo 2013; Beyene *et al.* 2015; Sleper and Bernardo 2018; Tiede and Smith 2018).

These models and selection methods implicitly assume that the breeding population has already been developed from selected parents. Therefore, the genetic variance of each trait, and the genetic correlation between traits, both of which determine the direct or correlated response to selection (Falconer and Mackay 1996), are fixed parameters of the population. In addition to more accurate selection within an established population, multi-trait genetic gain could be increased by developing better populations through deliberate selection of parent combinations with a more ideal mean, larger genetic variance, and more favorable genetic correlation. Typically, breeders select parents using the expected population mean, which can reliably be predicted as the mean of the two parents (Bernardo 2010). Recently developed methods that rely on *in silico* simulations (Bernardo 2014; Mohammadi *et al.* 2015) or deterministic equations (Zhong and Jannink 2007; Lehermeier *et al.* 2017; Osthushenrich *et al.* 2017) have been proposed to predict the genetic variance in a potential cross. These procedures model the expected segregation of genomewide markers with estimated effects, and early validation experiments suggest that such procedures may be useful (Lian *et al.* 2015; Tiede *et al.* 2015; Osthushenrich *et al.* 2017; Neyhart and Smith 2019).

Predictions of the population mean and genetic variance could be used to discriminate among potential crosses on the basis of the expected mean of selected progeny in those crosses. This can be quantified by the usefulness criterion (Schnell and Utz 1975), or the superior progeny mean (Zhong and Jannink 2007). The superior progeny mean assumes selection on a single trait, yet if two traits are genetically correlated, a response to selection would also be expected in a second trait. This “correlated progeny mean,” as we will refer to it, could be used to further distinguish ideal crosses as long as the genetic correlation is known or can be predicted. Though much research has focused on predicting the genetic variance in breeding crosses (e.g. Souza and Sorrells 1991; Bohn *et al.* 1999; Utz *et al.* 2001), little work has addressed predicting the genetic correlation. The simulation approach codified by Mohammadi *et al.* (2015) generates such predictions, but their accuracy and utility remain unexplored, and the use of simulations can be computationally burdensome for a large number of potential crosses.

The ideal selection of crosses to simultaneously improve multiple traits has been the focus of recent research. Allier *et al.* (2019) presented theory and a deterministic equation to predict the genetic correlation between two traits in multi- or bi-parental populations. While they applied this equation to the case of parental contribution (treated as a quantitative trait) correlated with an agronomic trait of interest, the theory could be generalized to two or more traits in the traditional sense. Additionally, Akdemir *et al.* (2019) applied a multi-objective optimized breeding strategy in simulations to select parent combinations and improve two unfavorably correlated traits. This approach solves the multiple objective optimization problem of maximizing the genetic gain of two or more traits while constraining inbreeding. Their maximization objective accounts for both the predicted mean and genetic variance of a cross, but does not consider the predicted genetic correlation between traits. In theory, such information could be included in this optimization framework, as long as predictions are accurate.

The objectives of this study were to (i) use simulations to assess the accuracy of genomewide predictions of genetic correlations and the long-term response to selection when selecting crosses on the basis of superior/correlated progeny means; and (ii) empirically measure the ability to predict genetic correlations using data from a barley (*Hordeum vulgare* L.) breeding program.

## METHODS AND MATERIALS

### Theory

Below, we first outline a deterministic prediction of the genetic variance of a single trait and the correlation between traits in a recombinant inbred line (RIL) population assuming two fully inbred parents, bi-allelic QTL, and no dominance or epistasis. This derivation follows the notation presented in Zhong and Jannink (2007); others have determined equations for the expected genetic variance in bi-parental populations of intermediate selfing generations (Lehermeier *et al.* 2017) or multi-parent populations (Allier *et al.* 2019), and this derivation could be applied to such circumstances. We then use these predictions to determine the superior progeny mean and correlated progeny mean for a cross.

Suppose that *L*_(*k*)_ QTL influence the *k*th quantitative trait, and in the *m*th cross *L*_*m*(*k*)_ QTL are segregating for that trait (where *L*_*m*(*k*)_ ≤ *L*_(*k*)_). The expected genetic variance in the cross is the sum of the variance of each locus plus the covariance between pairs of loci. As noted in Zhong and Jannink (2007), the genetic variance in cross *m* is

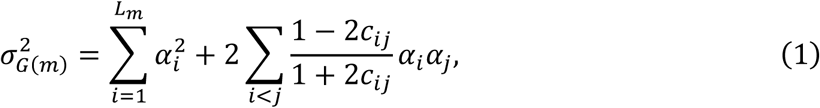

where *α*_*i*_ and *α*_*j*_ are the allele substitution effects of the *i*th and *j*th loci, respectively and *c*_*ij*_ is the recombination fraction between the *i*th and *j*th loci. As expected, loci that are genetically unlinked (i.e. independent, *c*_*ij*_ = 0.5) will have a covariance of 0. The covariance can be generalized across coupling and repulsion phase linkage by setting the allele substitution effects +*α*_*i*_ and +*α*_*j*_ to those of the first parent and −*α*_*i*_ and −*α*_*j*_ to those of the second parent (Zhong and Jannink 2007).

The single-trait covariance term in Equation (1) can be modified to calculate the expected genetic covariance between two traits, 1 and 2:

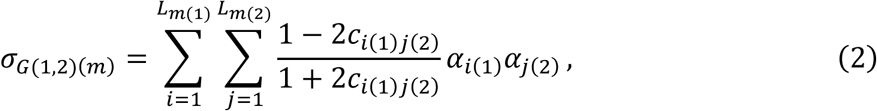

where *c*_*i*(1)*j*(2)_ is the recombination fraction between the *i*th locus of trait 1 and the *j*th locus of trait 2, *α*_*i*(1)_ is the allele substitution effect of the *i*th locus of trait 1 and *α*_*j*(2)_ is the allele substitution effect of the *j*th locus of trait 2. Using the expected genetic variance of each trait calculated from Equation (1) and the expected covariance between traits from Equation (2), the expected genetic correlation is

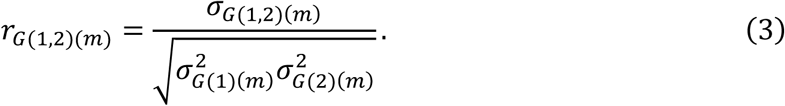

With estimates of the genetic variance for two traits and the genetic correlation between the traits, we can rely on established theory to estimate the superior progeny mean (*µ*_*sp*(*m*)_) and correlated progeny mean in a cross. For trait 1, assumed under direct selection, the superior progeny mean is

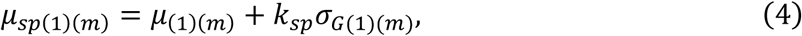

where *µ*_(1)(*m*)_ is the expected mean of trait 1 in the cross (estimated as the mean breeding value of the parents) and *k*_*sp*_ is the standardized selection coefficient. It is worth noting that the deviation from *µ*_(1)(*m*)_ in Equation (4) is the same as the direct response to selection, *R*_(1)_ = *k*_*sp*_*σ*_*G*(1)_, when the heritability is 1. The correlated response of the second trait, after selection on the first, is 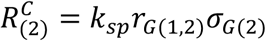, which, when expressed as a deviation from the expected mean of the second trait, becomes the correlated progeny mean:

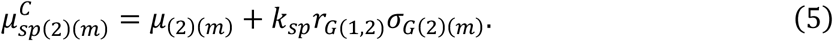

As with phenotypic values of two traits in a population, estimates of the superior progeny mean and correlated progeny mean could be used to select crosses that maximize the genetic gain for both traits, through independent culling levels or index selection (Bernardo 2010).

The equations above assume that the loci under consideration are the true QTL influencing the quantitative traits. Since the effects of such QTL are usually unknown, the estimated effects of genomewide markers in linkage disequilibrium with QTL can be used to make predictions (Meuwissen *et al.* 2001). This is the basis of *in silico* methods to predict genetic variance and genetic correlation, such as the R package *PopVar* (Mohammadi *et al.* 2015). The advantage of the deterministic equations is computational speed (about 130-fold faster, data not shown), with a high or perfect correlation between predicted values (Figure S1).

### Simulations

We conducted two simulations to assess the utility of predicting the genetic correlation in a breeding cross. Our simulations were based on observed marker genotypes of 1,570 North American two-row spring barley lines genotyped with 3,072 single nucleotide polymorphism (SNP) markers (Close *et al.* 2009), with genetic positions according to a consensus linkage map (Muñoz-Amatriaín *et al.* 2011); all data was obtained from the Triticeae Toolbox (https://triticeaetoolbox.org/barley/; Blake *et al.* 2016). We used empirical marker data to initiate our simulations in order to reflect the observed patterns of LD within North American barley breeding lines (Jannink 2010). Marker genotypes were arbitrarily coded as −1, 0, 1, where −1 was homozygous for the second allele, 0 was heterozygous, and 1 was homozygous for the first allele. After removing monomorphic and redundant SNPs (identical genotype calls and genetic positions), and SNPs and lines with more than 10% missing data, we were left with a marker matrix of 1,565 lines and 2,309 SNPs. We set the few heterozygous genotypes to missing and imputed missing calls using the mode across each SNP. The genetic map positions of completely coincident SNPs (i.e. due to low genetic resolution) were jittered by adding a small value (1 × 10^-6^ cM). These data were used to define the genetic architecture of the simulated quantitative traits and form the initial pool from which to establish a base training population.

#### Simulation 1 – Accuracy of predicting genetic correlations

In the first simulation experiment, we assessed the conditions influencing prediction accuracy of genetic correlations. We perturbed the heritabilities of two traits, the architecture defining the traits (i.e. number of QTL) and genetic correlation, the initial genetic correlation, and the base/training population size (Table 1). Simulations were initiated by drawing 200 SNP markers to act as QTL. For each trait, 100 – *L* QTL were assigned an effect of 0, where *L* was the effective number of QTL (30 or 100). QTL effects were defined by a geometric series, as proposed by Lande and Thompson (1990): for the *k*th QTL, the value of the favorable homozygote was *a*^*k*^, the value of the heterozygote was 0, and the value of the unfavorable homozygote was −*a*^*k*^, where *a* = (1 − *L*)/(1 + *L*). The first allele at each QTL was randomly assigned to be favorable or unfavorable and larger values were considered favorable for both traits. This randomization was performed independently for each trait.

**Table 1.** In our two simulation experiments, we modified the heritability (*h*^*2*^) and number of quantitative trait loci (*N*_*QTL*_) of two traits, the starting genetic correlation (*r*_*G(0)*_), correlation architecture, size of a training population (*N*_*TP*_), and model used to predicted genomewide marker effects.

| Simulation experiment | $h^2$ | $N_{QTL}$ | $r_{G(0)}$ | Correlation architecture | $N_{TP}$ | Model |
| --- | --- | --- | --- | --- | --- | --- |
| Experiment 1<br>(prediction accuracy) | Trait 1: 0.3, 0.6, 1<br>Trait 2: 0.3, 0.6, 1 | 30, 100 | -0.5, 0, 0.5 | Pleiotropy<br>Tight linkage<br>Loose linkage | 150, 300,<br>450, 600 | RR-BLUP<br>BayesCπ |
| Experiment 2<br>(long-term response) | Trait 1: 0.6<br>Trait 2: 0.3, 0.6 | 100 | -0.5, 0.5 | Pleiotropy<br>Tight linkage<br>Loose linkage | 600 | RR-BLUP |

Genetic correlations were generated according to three different architecture types: pleiotropy, tight linkage, or loose linkage. For simplicity, we assumed that the genetic architecture was governed entirely by one of the types. Under pleiotropy, the sampled QTL effects were first stored in an *L* × 2 matrix, **A**. The desired genetic correlation in the base population (*r*_*G*(0)_) was achieved by multiplying matrix **A** by the Choleski decomposition of the variance-covariance matrix **Σ**, which contained 1 on the diagonal and *r*_*G*(0)_ on the off-diagonal. This resulted in a set of QTL with pleiotropic effects that varied in both magnitude and sign for the two traits. Under tight linkage and loose linkage, each SNP sampled to be an effective QTL for the first trait was paired with another SNP that was sampled – with restrictions – to be an effective QTL for the second trait. For tight linkage, this second SNP was restricted to within 5 cM of the first SNP, and for loose linkage, this second SNP was restricted to between 25 cM and 35 cM of the first SNP. The QTL effects were again stored in the *L* × 2 matrix **A**, where each row was a pair of QTL, and subsequently adjusted as above. Effects of QTL influencing the second trait were then multiplied by matrix **R**, which contained estimates of linkage disequilibrium (measured as the pairwise correlation, *r*, between genotype states in the base population) between QTL influencing the first trait and QTL influencing the second trait. This adjustment resulted in base genetic correlations that approximately matched the target, *r*_*G*(0)_ (Figure S2).

The base/training population was first generated by randomly sampling *N*_*TP*_ individuals from the simulation starting material. For each trait, the genotypic value of an individual was calculated as the sum of the QTL allele effects carried by that individual, and the genetic variance was calculated as the variance among genotypic values. Phenotypic values were simulated by adding independent normally distributed deviations to the genotypic values to achieve a starting entry-mean heritability of 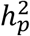 (Table 1) with no residual covariance between traits. Individuals were assumed to be phenotyped in three environments with one replication, and the mean phenotypic value was used for genomewide prediction. Marker effects were predicted using the univariate model:

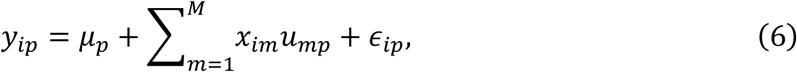

where *y*_*ip*_ was the phenotypic mean of the *i*th individual for the *p*th trait, *µ*_*p*_ was the population mean for the *p*th trait, *x*_*im*_ was the allelic state of the *m*th marker in the *i*th individual (i.e. −1, 0, or 1), *u*_*mp*_ was the predicted effect of the *m*th marker for the *p*th trait, and *ϵ*_*ip*_ was the associated error. We used two models to predict marker effects: ridge-regression best linear unbiased prediction (RR-BLUP) and BayesCπ (Habier *et al.* 2011). Potential crosses were generated by randomly sampling 50 pairs of individuals from the base population. We predicted the genetic correlation 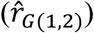 for each potential cross using Equations (1), (2), and (3), where *α* was substituted with *u*_*mp*_. The expected genetic correlation (*r*_*G*(1,2)_) was similarly computed but instead using the known QTL effects instead of the predicted marker effects. Prediction accuracy was defined as the correlation between the predicted and expected genetic correlations. As a comparison, we also assessed predictions of the trait-specific mean 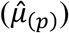 and genetic variance 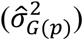 in each cross. Each condition of this simulation was replicated 100 times.

#### Simulation 2 – Correlated response to selection

We conducted a second simulation experiment to measure the long-term response of two correlated traits under different cross selection strategies. The range of perturbed parameters was smaller than in the first simulation (Table 1), though simulations were initiated as described above. We assumed that a breeder wanted to simultaneously increase the genotypic value of two quantitative traits through indirect selection, therefore positive genetic correlations were favorable. The base/training population was created by randomly sampling *N*_*TP*_ = 600 individuals from the simulation starting material. Informed by the results of the first simulation (see below), we used only the simpler RR-BLUP model to predict marker effects. Potential parents for the first breeding cycle were selected by determining the best 30 individuals in training population based on the predicted genotypic values of the first trait (the primary trait under direct selection). We predicted the mean, genetic variance, and genetic correlation for all possible non-reciprocal crosses between the potential parents. We then calculated the superior progeny mean of the first trait and the correlated progeny mean of the second trait using the predicted parameters and Equations (4) and (5) with a standardized selection coefficient of *k*_*sp*_ = 2.06 (i.e. selection of the best 5%).

Twenty crosses were selected based on i) equal-weight sum of the normalized predicted superior progeny mean of the first trait and predicted correlated response in the second trait 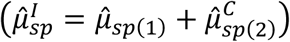, ii) the predicted cross mean of the primary trait 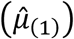, or iii) random selection. We will subsequently refer to the non-random cross selection methods by the abbreviations CPM (correlated/superior progeny mean) or FM (family, or cross, mean). From the selected crosses, families of 50 recombinant inbred lines were simulated using the *qtl* R package (Broman *et al.* 2003). Recombination events were sampled according to the genetic map (Muñoz-Amatriaín *et al.* 2011), with the assumption of no crossover interference or mutation. This resulted in a pool of 1,000 selection candidates. Finally, 50 potential parents for the next breeding cycle were chosen from these candidates using predicted genotypic values of the first trait. We simulated 10 cycles of recurrent selection (outlined in Figure 1), during which marker effect estimates remained unchanged. Along with the standardized selection response for each trait, we also tracked the genetic variance of each trait, the genetic correlation between traits, the frequency of favorable, unfavorable, and antagonistic (i.e. two alleles with opposite effect) QTL haplotypes, and the proportion of QTL with fixed alleles. Simulations were replicated 250 times, and we report the mean and 95% approximate confidence interval for each measured variable.

**Figure 1.**
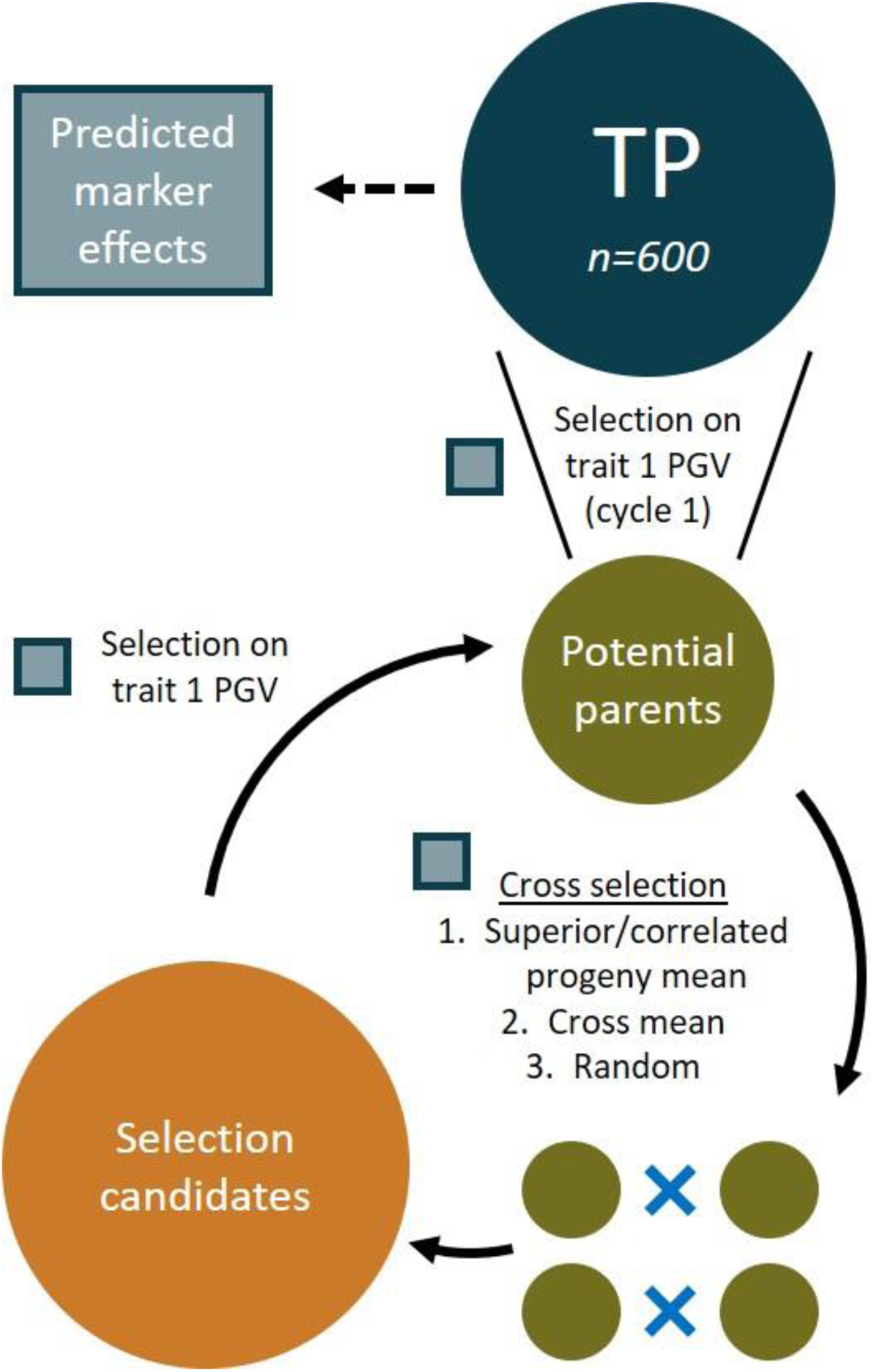
In the recurrent selection simulation (Simulation 2), a training population (TP) was sampled and used to predict genomewide marker effects. In the first cycle, potential parents were identified using direct selection on predicted genotypic values (PGVs) of the first trait. Crosses were selected by one of three methods and were used to simulate selection candidates. Potential parents of the next cycle were determined using direct selection on the first trait. Ten breeding cycles were simulated. Any processes that relied on the predicted marker effects are noted with a blue/grey box.

### Empirical validation

To empirically validate predictions of genetic correlations, we used phenotypic and genotypic data from a barley breeding program. The details of data generation are described elsewhere (Neyhart and Smith 2019), but we include a brief overview below. A training population (TP) of 175 two-row spring barley lines was genotyped with 6,361 SNP markers and phenotyped in four location-year environments for heading date (a proxy for flowering time), Fusarium head blight (FHB) severity (caused by the fungal pathogen *Fusarium graminearum* Schwabe), and plant height. To estimate the genetic correlation between pairs of traits in the TP, we regressed the phenotypic observations of two traits, ***y***_*p*_ = {*y*_*ijp*_}, on individuals (genotypes), environments, and their interaction, where *i* indexes genotypes, *j* environments, and *p* traits. We fitted a bi-variate mixed model:

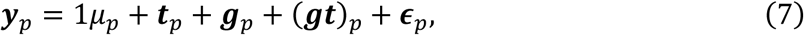

where *µ*_*p*_ is the population mean, ***t***_*p*_ = {*t*_*jp*_} the fixed environmental effect, ***g***_*p*_ = {*g*_*ip*_} the random effect of genotypes, (***gt***)_*p*_ = {(*gt*)_*ijp*_} the random interaction effect of genotypes and environment, and ***ϵ***_*p*_ = {*ϵ*_*ijp*_} the associated error. The distribution of random effects was assumed to be multivariate normal such that: 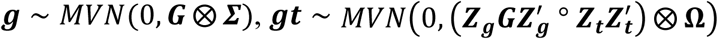, and ***ϵ*** ∼ *MVN*(0, ***I*** ⨂ ***R***), where ***G*** is the realized genomic relationship matrix, ***Z***_***g***_ is an incidence matrix for individuals, ***Z***_***t***_ is an incidence matrix for environments, ***I*** is an identity matrix, ⨂ denotes the Kronecker product between matrices and ° denotes the Hadamard product between matrices. The genotype, genotype-environment interaction, and residuals covariance structures are

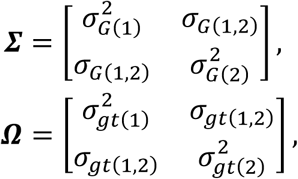

and

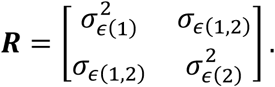

Models were fitted using the R package *sommer* (Covarrubias-Pazaran 2016) and the genetic correlation was estimated from elements in ***Σ*** using Equation (3).

Marker effects were estimated from marker genotypes and phenotypic best linear unbiased estimates (BLUES; i.e. genotypic means from a model accounting for genotype, environment, and the interaction) of the TP using the univariate model in Equation (6). Among all potential non-reciprocal crosses between 813 offspring of the TP (*n* = 330,078), we used estimated marker effects and the R package *PopVar* to predict the genetic correlation for each pair of traits. (Predictions were generated early in the study using this package, so for consistency we used those values, and not those generated using the deterministic equations above.) Twenty-six crosses were made based on the predictions, producing “validation families” ranging from 28 to 160 F_5_ lines. The criteria for selecting crosses are described in Neyhart and Smith (2019) and rested primarily on predictions of genetic variance for several traits relevant for the breeding program. Validation families were phenotyped for the same three traits in 2 or 4 environments. Observations of heading date and plant height were recorded for all families, but due to logistical constraints of the inoculated disease nursery, only 14 families were phenotyped for FHB severity.

Validation family phenotypes were used to estimate the observed genetic correlation in each family. We fitted a model modified from Equation (7):

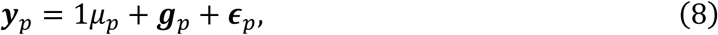

where ***y***_*p*_ = {*y*_*ip*_} was the BLUE of genotypes in a validation family and other terms as described above. The distribution of random genotypic effects was assumed multivariate normal such that ***g*** ∼ *MVN*(0, ***I*** ⨂ ***Σ***), where the genomic relationship matrix in Equation (7) was replaced by an identity matrix, since validation families were ungenotyped equally related within a family. Predictive ability was measured as the correlation between predicted and estimated genetic correlations across validation families, and the significance of this coefficient was tested using 1,000 bootstrapping replicates. Note that predictive ability is calculated by comparing predictions with phenotype-based observations, whereas prediction accuracy compares predictions with the true genotypic parameter (unobservable in our empirical experiment).

### Data availability

Marker data for initiating the simulations and all data used in the empirical validation experiment is available from the Triticeae Toolbox (T3; https://triticeaetoolbox.org/barley/). All simulations and analyses were performed in R (v. 3.5.1; R Core Team, 2018) and relevant scripts are located in the GitHub repository https://github.com/UMN-BarleyOatSilphium/GenCorPrediction.

Instructions are included in this repository for downloading data from T3. Supplementary figures and tables are available through Figshare.

## RESULTS

### Factors influencing predictions of genetic correlation

In our simulation, prediction accuracy for the genetic correlation in potential crosses was most influenced by trait heritability, training population size (*N*_*TP*_), and genetic architecture. We provide a cross-section of results in Figure 2, and all results for the first simulation are displayed in Figure S3 and Table S1. Accuracy increased additively as a function of the heritability of both simulated traits, but only reached a maximum of about 0.81 under the most ideal conditions (Figure S3, Table S1). On average, accuracy increased by about 1.5-fold when moving from *N*_*TP*_ = 150 to *N*_*TP*_ = 600. We generally did not observe a pattern of diminishing returns when increasing *N*_*TP*_, though some evidence of that pattern was present under the tight linkage architecture (Figure 2).

**Figure 2.**
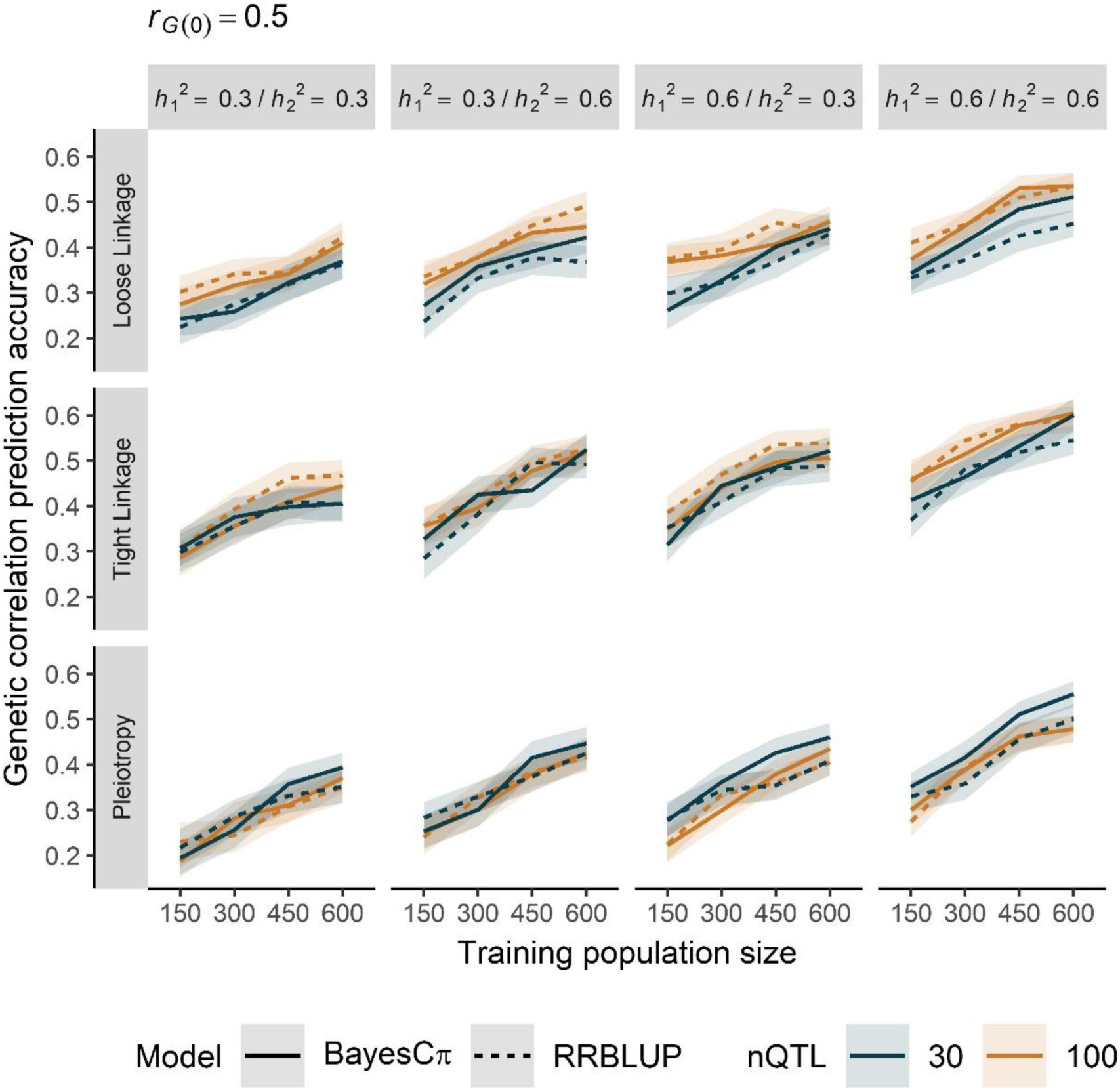
The prediction accuracy of the genetic correlation in a cross increased with larger training populations and greater trait heritability. Accuracy was also influenced by genetic correlation architecture (loose linkage, tight linkage, or pleiotropy), prediction model [BayesCπ (solid) or RRBLUP (dashed)], and number of QTL [30 (navy) or 100 (orange)]. Lines denote the mean of 100 simulation replicates, and ribbons denote a 95% confidence interval. Results are restricted to a base genetic correlation of 0.5. (See Figure S3 and Table S1 for complete results.)

Between all correlation architectures, tight linkage resulted in the highest prediction accuracy, followed by loose linkage and then pleiotropy. The difference in accuracy under tight linkage versus loose linkage was on average 0.071 (17%) and this difference under loose linkage versus pleiotropy was 0.017 (6.2%). With tight linkage and loose linkage genetic architectures, accuracy was slightly higher when 100 versus 30 QTL influenced both traits (a difference of about 0.04, or 10%), but the reverse was true under pleiotropy, where the accuracy was about 8% lower (a difference of about 0.03) with more QTL (Figure 2, Table S1). Interestingly, an interaction was apparent between the genetic architecture and the prediction model. Under pleiotropy and tight linkage, there was a slight advantage to using the BayesCπ model over RR-BLUP, particularly when 30 QTL influenced each trait. This difference was quite slim, however, with a boost to accuracy of only about 0.015 (3%).

The family mean and genetic variance of each trait in potential crosses were almost always predicted with greater accuracy than the genetic correlation (Figure 3, Table S1). In general, the family mean was predicted most accurately, followed by the genetic variance and the genetic correlation. (The genetic variance of one trait was occasionally predicted more accurately than the family mean of another trait, but only if the heritability of the first trait was much less than the second.) This trend was consistent across training population sizes, prediction models, and genetic architectures. The average (and range in) prediction accuracy was 0.87 (0.64, 1.0) for the family mean, 0.66 (0.34, 0.96) for the genetic variance, and 0.48 (0.18, 0.81) for the genetic correlation.

**Figure 3.**
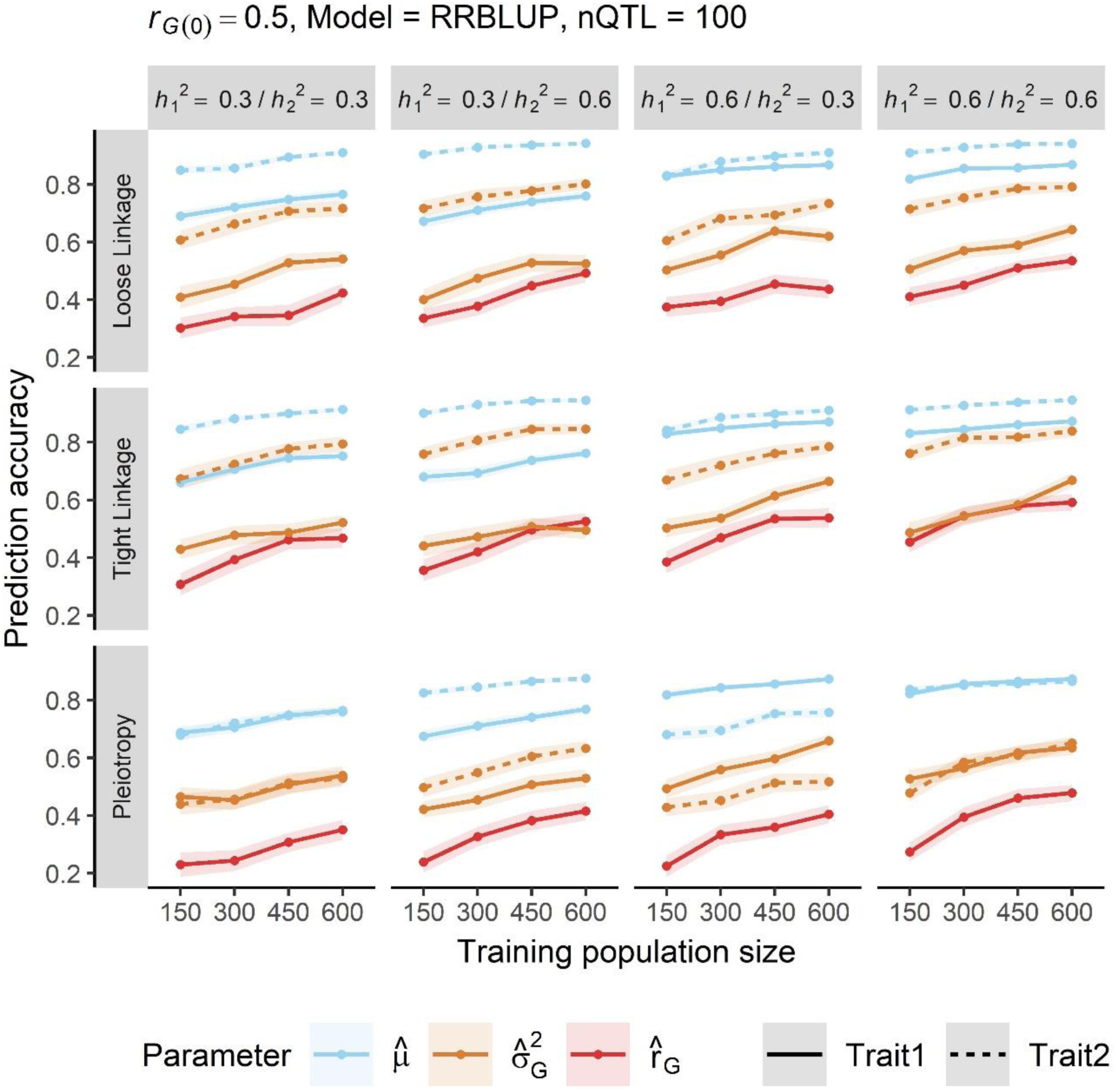
Parameters of a cross were predicted with varying degrees of accuracy in our simulation. Predictions of the cross mean (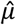, blue) were most accurate, followed by genetic variance (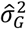, orange) and genetic correlation (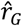, red); this ranking was consistent across trait heritabilities, training population size, and genetic correlation architecture (loose linkage, tight linkage, or pleiotropy). Lines denote the mean of 100 simulation replicates, and ribbons denote a 95% confidence interval. Results are restricted to a base genetic correlation of 0.5, RRBLUP prediction model, and 100 QTL. (See Table S1 for complete results.)

### Empirical validation of predicted genetic correlations

We used genomewide markers and phenotypic data to empirically estimate the genetic correlation for three pairs of quantitative traits in our 175-line training population (TP). The genetic correlation was −0.84 between *Fusarium* head blight (FHB) severity and heading date, - 0.44 between FHB severity and plant height, and 0.48 between heading date and plant height (Table 2). These estimates were reflected in predictions of the cross mean and genetic correlation of 330,078 potential crosses (Figure 4, Table 2). The average predicted genetic correlation among the potential crosses was −0.54 for FHB severity and heading date, −0.26 for FHB severity and plant height, and 0.24 for heading date and plant height. Though the genetic correlations between FHB severity and both heading date and plant height were unfavorable (earlier-flowering, shorter, and disease resistant plants are desirable), predictions implied that progress could be made by selecting populations with more favorable genetic correlations. For instance, more than 2,400 (0.73%) potential crosses were predicted to have a favorable (i.e. positive) correlation between FHB severity and heading date.

**Table 2.** Estimated genetic correlation in the empirical barley training population 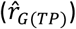 and the mean and range of the predicted genetic correlations among 330,078 potential crosses 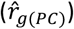 for three pairs of quantitative traits

| Trait 1 | Trait 2 | $\hat{r}_{G(TP)}$ | Mean (range) of $\hat{r}_{g(PC)}$ |
| --- | --- | --- | --- |
| FHB<sup>a</sup> Severity | Heading Date | -0.84 | -0.54 (-0.94, 0.52) |
| FHB Severity | Plant Height | -0.44 | -0.26 (-0.86, 0.63) |
| Heading Date | Plant Height | 0.48 | 0.24 (-0.73, 0.87) |

<sup>a</sup>Fusarium head blight

**Figure 4.**
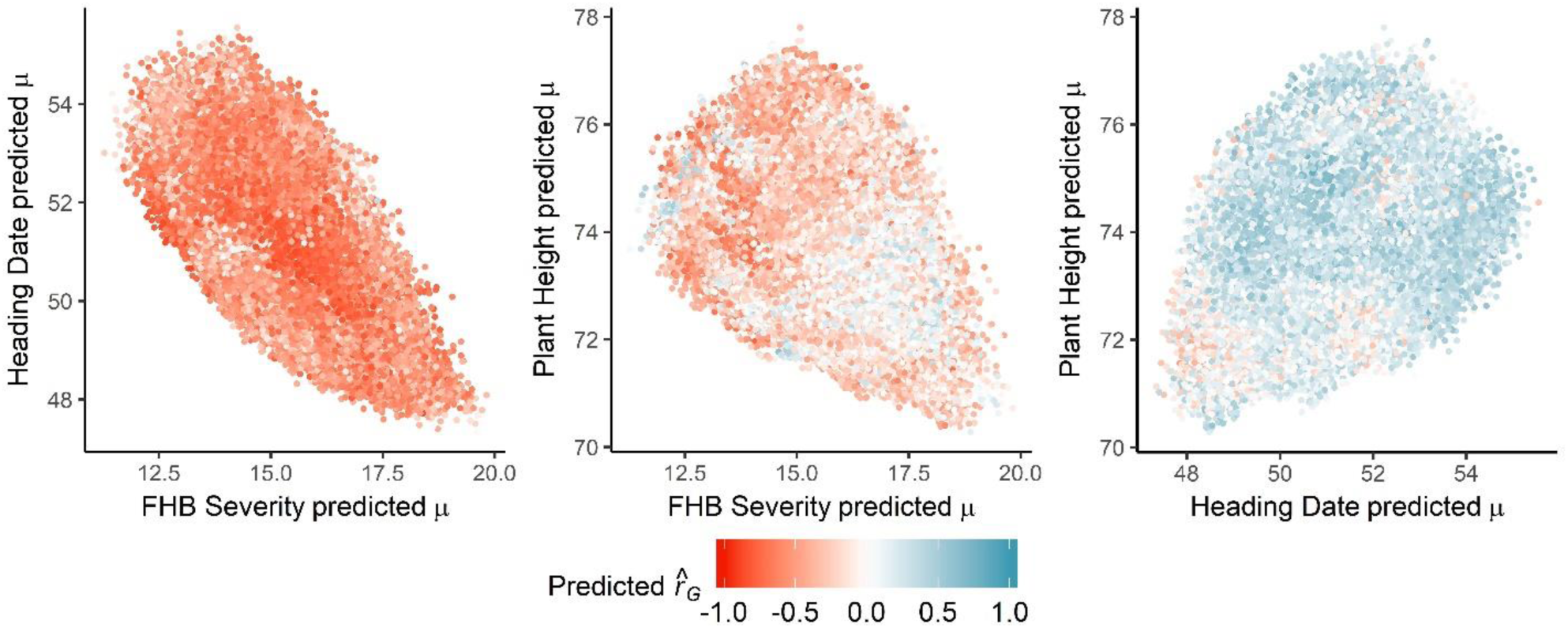
In the barley breeding population used to empirically validate predictions, the relationship between the predicted cross means 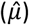 of three pairs of traits for *n* = 330,078 potential crosses mirrored the overall distribution of predicted genetic correlations 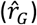, shaded from red (negative) to blue (positive). FHB, Fusarium head blight.

The mean (and range) of estimated genetic correlations among the validation families was −0.18 (−0.72, 0.58) for FHB severity and heading date, −0.038 (−0.67, 0.64) for FHB severity and plant height, and −0.13 (−0.64, 0.69) for heading date and plant height (Table 3). Estimates of predictive ability for genetic correlations ranged from −0.012 to 0.41 (Table 3). We could only validate predictions of the correlation between heading date and plant height, where all 26 validation families (VF) were phenotyped. The predictive abilities for remaining trait combinations were not significantly different from zero (*P* > 0.05; bootstrapping). The ability to predict the genetic correlation appeared to coincide with the heritability of both traits; the entry-mean heritability in the TP (and in the VF) was 0.45 (0.11) for FHB severity, 0.96 (0.78) for heading date, and 0.52 (0.74) for plant height (Neyhart and Smith 2019).

**Table 3.** Number of phenotyped validation families, mean (and range) of observed genetic correlations, and the predictive ability, measured as the correlation between the predicted 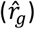 and observed (*r*_*G*_) genetic correlation, for each of three pairs of traits. A 95% confidence interval for the predictive ability was estimated from 1,000 bootstrapping replicates.

| Trait 1 | Trait 2 | $N_f^b$ | Mean (range) of $r_g$ | $cor(\hat{r}_g, r_g)$ |
| --- | --- | --- | --- | --- |
| FHB<sup>a</sup> Severity | Heading Date | 14 | -0.18 (-0.72, 0.58) | 0.24 (-0.30, 0.70) |
| FHB Severity | Plant Height | 14 | -0.038 (-0.67, 0.64) | -0.012 (-0.53, 0.59) |
| Heading Date | Plant Height | 26 | -0.13 (-0.64, 0.69) | 0.41* (0.024, 0.71) |

<sup>a</sup>Fusarium head blight
<sup>b</sup>Number of families used to compute predictive ability
\*Significant at $P < 0.05$

### Long-term response with different cross selection strategies

Our second simulation showed that the genetic gain for two correlated quantitative traits was impacted by the base genetic correlation, the genetic architecture, and the strategy to select crosses (Figure 5). We found little difference in the outcome when the heritability of the second trait was 0.6; therefore, we highlight results when the heritability of the second trait was 0.3, a more realistic situation for indirect selection (Bernardo 2010). When measuring progress via a trait index (Figure 5A), we found that selecting crosses based on the predicted correlated superior progeny mean (CPM) resulted in a greater response than selection on the predicted cross mean (FM) or by random selection. Under all genetic architectures, the advantage of imposing non-random cross selection became clear after 1 breeding cycle. Subsequently, after 3 cycles, selecting crosses based on CPM resulting in higher gain than by selecting based on FM. Only after 9 – 10 cycles did random cross selection achieve equivalent or superior genetic gain compared with FM selection, though it never outperformed selection using CPM.

**Figure 5.**
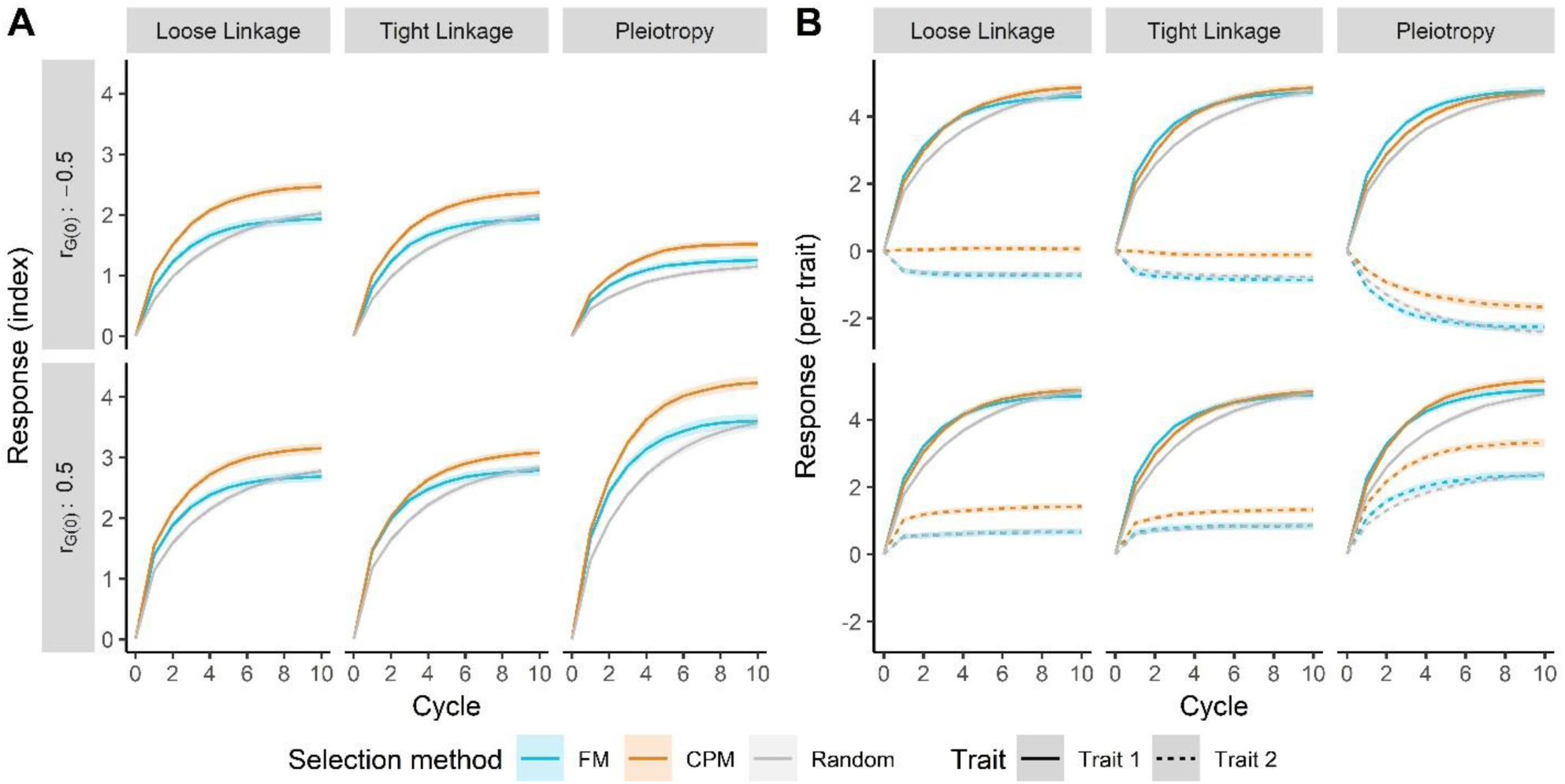
Selecting crosses on the predicted correlated/superior progeny mean (CPM, orange) led to a greater long-term response (in units of genetic standard deviations) compared to selection on the predicted family mean (FM), blue) or by random selection (grey). This was true for a two-trait index (**A**) and both traits individually (**B**) across three correlation architectures and two base genetic correlations (*r*_*G(0)*_). Lines denote the mean of 250 simulation replicates, and the ribbon denotes a 95% confidence interval.

Gain from selection, and marginal differences between selection methods, depended on the genetic architecture and correlation. The final genetic gain was, on average, less when the genetic correlation was negative. With pleiotropic architecture, the reduction in genetic gain from a correlation of 0.5 to −0.5 was more severe (a roughly 100% decrease) than with tight or loose linkage architectures (a roughly 20% decrease). After 10 cycles, the marginal genetic response (based on an index) when using CPM versus FM cross selection ranged from 0.30 (11%), with tight linkage architecture and positive correlation, to 0.53 (27%), with loose linkage architecture and negative correlation.

When considering traits individually, we found that much of the advantage of selecting crosses on CPM was realized in the correlated response of the second trait (Figure 5B). Selection on CPM or FM yielded similar responses in the first trait, except under pleiotropic architecture and a positive genetic correlation. Again, random cross selection led to the lowest response, though in some cases it became equivalent to selection on CPM or FM after 10 breeding cycles. Genetic gain was consistent for the primary trait, with a plateau reached after 6 – 8 cycles of selection. Conversely, we observed a rapid plateauing in the genetic gain of the second trait when the architecture was not pleiotropic. Here, when the genetic correlation was positive, the correlated response in the second trait was also positive, as expected, with a greater response under CPM selection. When the genetic correlation was negative, FM or random cross selection led to a negative response in the second trait, which was prevented under CPM selection (Figure 5B). Under pleiotropic conditions, the correlated response in the second trait matched expectations given the genetic correlation; however, CPM selection increased the positive response with positive correlation and mitigated the negative response with negative correlation.

As expected, the genetic variance for both traits decreased over cycles of selection (Figure 6A), and by cycle 10, most had been exhausted. Although genetic variance for the first trait declined similarly under varying architectures, the loss of variance for the second trait was more precipitous when the architecture was defined by linkage versus pleiotropy. Under the latter, genetic variance for the second trait was reduced at a rate comparable to the first trait (Figure 6A). We found that cross selection by FM always led to the most rapid reduction of genetic variance, while this decay was slower when selecting on CPM and slower yet with random mating. This ranking among selection methods was very apparent for the first trait (and for the second trait under pleiotropic architecture), but marginal differences were much less for the second trait.

**Figure 6.**
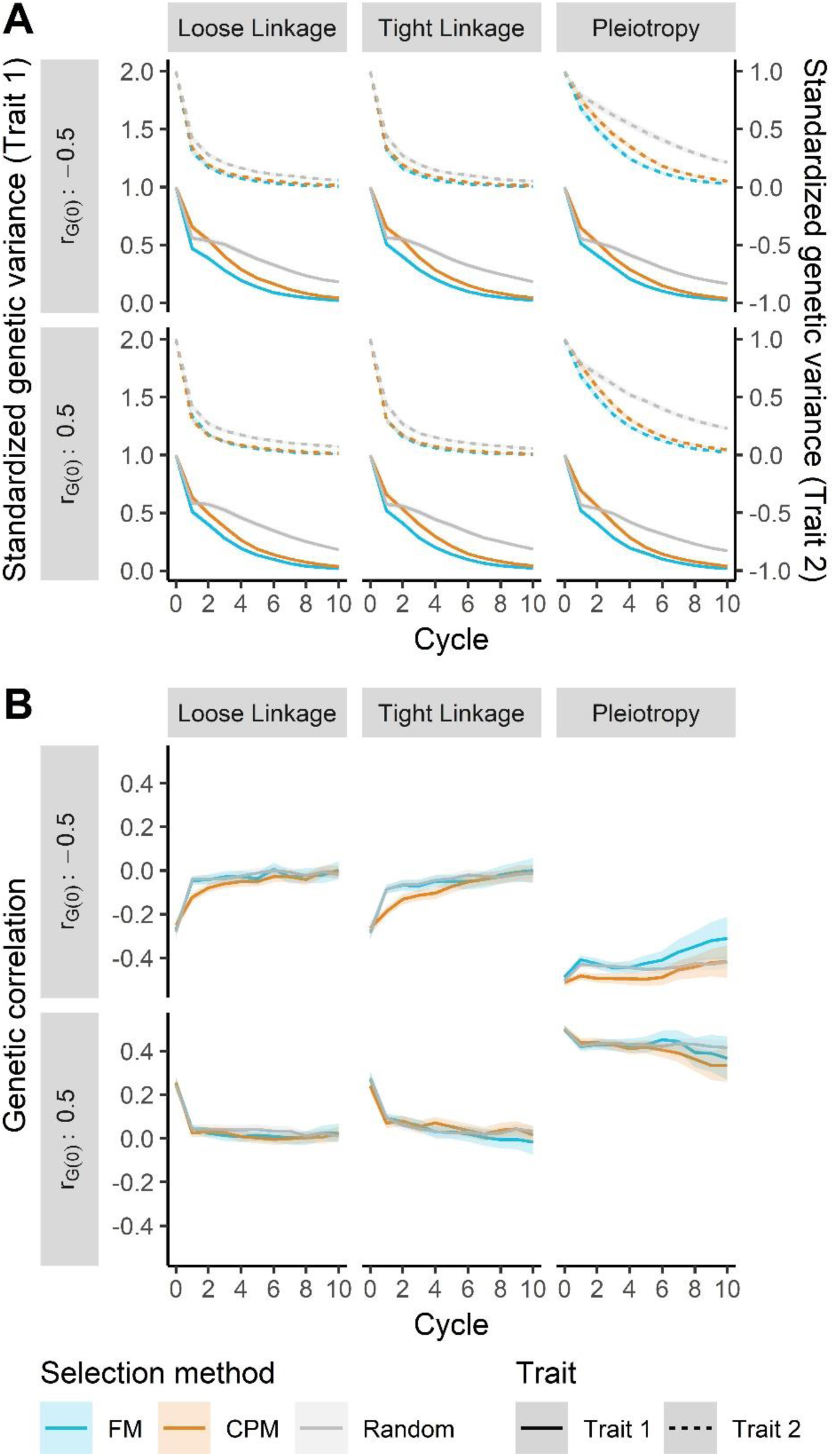
Genetic variances (**A**) and the genetic correlation (**B**) in the simulated indirect selection breeding program were influenced by the cross selection method (family mean, FM, blue; correlated/superior progeny mean, CPM, orange; random, grey), genetic architecture, and base genetic correlation (*r*_*G(0)*_). All components declined in absolute value over cycles of selection, but the rate depended on the cross selection method and genetic architecture. Lines denote the mean of 250 simulation replicates, and the ribbon denotes a 95% confidence interval.

The genetic correlation in the breeding population consistently declined in absolute value, moving towards zero under all simulated conditions (Figure 6B). The genetic architecture impacted the rate of change, with the most rapid movement under loose linkage, followed by tight linkage and then pleiotropy, as expected. The correlation initially became more negative when selection was imposed on the base population (i.e. cycle 0 to cycle 1). This change was much larger when the base correlation (*r*_*G*(0)_) was positive; indeed, under non-pleiotropic architecture, the genetic correlation became near-zero, or negative, after 1 cycle of selection. Conversely, when *r*_*G*(0)_ was negative, the genetic correlation moved more steadily towards 0. When *r*_*G*(0)_ was negative, we found that selecting crosses on CPM usually led to a more negative genetic correlation than FM or random selection, particularly in the first 5 breeding cycles.

Changes in haplotype frequency were greatest when loose linkage defined the correlation architecture, followed by tight linkage and pleiotropy (Figure 7A). The change in haplotype frequencies was more limited with negative *r*_*G*(0)_, particularly under pleiotropic architecture. Selecting crosses on CPM led to a significantly higher increase in the frequency of favorable haplotypes and a corresponding decrease in the frequency of unfavorable haplotypes. Further, with non-pleiotropic architecture we observed a greater reduction in the frequency of antagonistic haplotypes when selecting crosses by CPM than other methods. As expected, the frequency of antagonistic haplotypes did not change when the architecture was defined by pleiotropy (Figure 7A). Selection increasingly drove QTL to fixation (Figure 7B), but the rate of fixation was uneven for different cross selection methods. Choosing crosses on FM led to the highest fixation rate, followed by CPM and then random mating. There was a slightly higher fixation rate with positive genetic correlation than with negative genetic correlation, and the fixation rates for QTL influencing the first trait or second trait were roughly equivalent.

**Figure 7.**
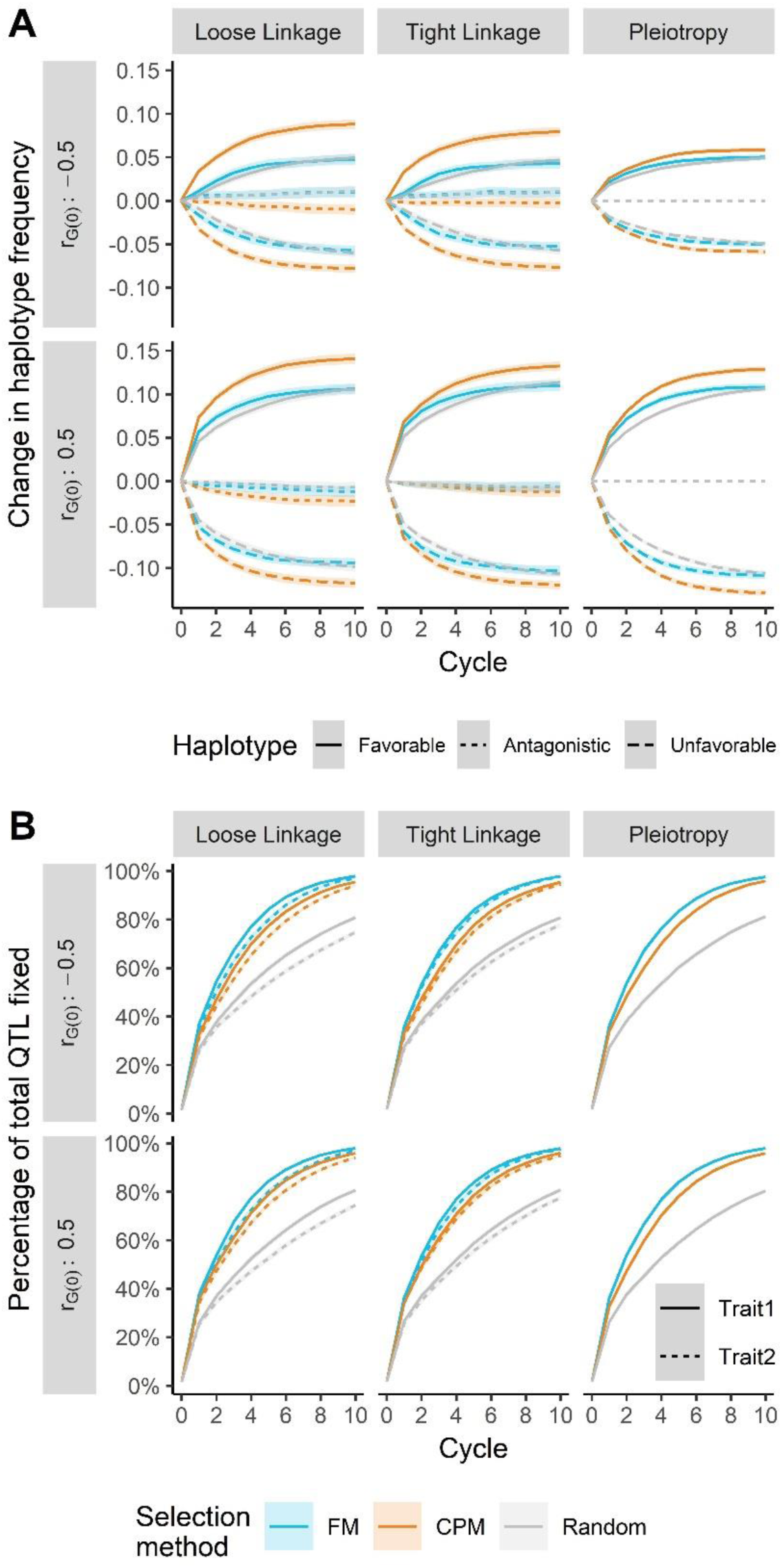
The change in frequency of two-trait quantitative trait locus (QTL) haplotypes and proportion of fixed QTL depended on the cross selection method genetic architecture, and base genetic correlation (*r*_*G(0)*_). (**A**) The increase in frequency of favorable (solid) haplotypes and decrease in frequency of unfavorable (dashed) and antagonistic (dotted) haplotypes was always greater when selecting crosses on the correlated/superior progeny mean (CPM, orange). (**B**) The rate of QTL fixation for both trait 1 (solid) and trait 2 (dotted) was always greatest with cross selection on the family mean (FM, blue), followed by CPM and random mating (grey). Lines denote the mean of 250 simulation replicates, and the ribbon denotes a 95% confidence interval.

## DISCUSSION

### Predictions of genetic correlations are feasible with reliable training data

Our first simulation measured the prediction accuracy of genetic correlations as a function of training population (TP) size, trait heritability, prediction model, and genetic architecture. Our results implicated the usual suspects driving genomewide prediction accuracy for individual traits. Increasing the TP size and the heritability of both traits improved accuracy, an expected result given the importance of these parameters (Daetwyler *et al.* 2008; Wimmer *et al.* 2013). The impact of genetic architecture was curious. Genetic correlations caused by loose linkage or pleiotropy led to lower prediction accuracies compared to architecture defined by tight linkage (Figure 2). We hypothesize that the same phenomenon, albeit with opposite effect, is responsible. Genomewide prediction models (i.e. RRBLUP or BayesCπ) assume that many more markers than true QTL have non-zero effect on both traits. Our approach to predicting genetic correlations relies on the recombination and segregation of these markers, implying that all contribute to variability in genetic variance and covariance, while only true QTL generate this variability. With pleiotropy, this would manifest as a downward bias in the covariance between traits, as we predict a greater possibility of recombination between markers than what is possible for the true QTL. Under loose linkage, an opposite, upward bias in covariance would be expected, as the many markers with uneven trait effects would be predicted to co-segregate more often than what is possible for the true QTL. Indeed, when we calculated the average bias of the predicted genetic correlations, we observed a roughly 30% upward bias under loose linkage and an opposite 30% downward bias under pleiotropy (Figure S4). Additionally, though the bias in predicting genetic covariance was always negative, it was less so under loose linkage than under pleiotropy. This bias, particularly if uneven across predicted crosses, could lead to the observed loss in accuracy. Practically, the impact of genetic architecture may be less important, since architecture is generally immutable and the effect on prediction accuracy is small (Figure 2). Though prediction accuracies were not appreciably different between models, it is worth mentioning potential causes and impacts. With fewer QTL, we observed higher prediction accuracies under the BayesCπ model, an unsurprising result given the known advantage of such models with heterogenous contributions of SNPs to total variance (Daetwyler *et al.* 2010); however, as expected, differences between models were smaller with architectures defined by more QTL. Although not observed in our simulations, we might expect the genetic diversity in the training population to play a role in prediction accuracy. Lower diversity, defined by lower effective population sizes and fewer distinct haplotype blocks, will lead to a spreading of the variance across few sets of markers in high LD (Daetwyler *et al.* 2008, 2010). This spreading would also exacerbate the observed upward bias under loose linkage architecture and downward bias under pleiotropic architecture. Further, the larger haplotype blocks accompanying lower diversity would cloud differences in predictions between tight linkage and loose linkage architectures. Predictions of genetic correlations in barley and other inbreeding crops may be particularly sensitive to this problem, given the long-range persistence of LD observed in these species (Hamblin *et al.* 2010; Chao *et al.* 2011).

Differences in the accuracy to predict the three parameters of a potential cross (i.e. mean, genetic variance, and genetic correlation) are attributable to the nature of each statistic and have practical implications. Greater accuracy when predicting the cross mean versus genetic variance was predicted by theory (Zhong and Jannink 2007) and has been observed empirically (Adeyemo and Bernardo 2019; Neyhart and Smith 2019); this trend is expected because the genetic variance, a second-order statistic, will be more adversely impacted by error in marker effect estimates. Similarly, the accuracy of the genetic correlation, a ratio of second-order statistics with large sampling variance (Robertson 1959), will be even more adversely affected. Even at large TP sizes, the predictions of the genetic correlation were only as accurate as those of the genetic variance at modest TP size and never as accurate as those of the cross mean (Figure 3). Practically, this suggests that very large TPs are needed for such predictions to be useful, a prospect that may be prohibitive for a plant breeder. Further research could be directed towards applying Bayesian approaches, such as the “posterior mean variance” method of Lehermeier *et al.* (2017), to improve the accuracy and bias of predicted genetic correlations, particularly at small TP sizes.

### Selecting crosses using predicted genetic correlations increases multi-trait response

Under a simulated breeding program focused on multi-trait recurrent indirect selection, we showed that the long-term genetic gain for two traits was greatest when crosses were selected on predicted correlated/superior progeny means (CPM). This was true under all conditions of genetic architecture, trait heritability, and base genetic correlation (*r*_*G*(0)_) (Figure 5). Further, cross selection based on CPM was superior to selection on the predicted cross mean (FM) or random mating, standard choices in programs using “best-by-best” breeding for cultivar development or in recurrent selection (Bernardo 2010).

The greater multi-trait selection response achieved under CPM cross selection can be attributed to many drivers, including genetic variance and correlation, haplotype and allele frequencies, and linkage disequilibrium (LD). We discuss their impacts below. Compared to FM, selection on CPM led to a higher maintenance of genetic variance for both traits (Figure 6A). When selecting on CPM, particularly at the relatively high selection intensity used in our simulation (*i* = 0.05; *k*_*sp*_ = 2.06), more weight is given to the predicted genetic variance versus the predicted mean (Equations 4 and 5). Segregation of QTL is explicitly driving the prediction of genetic variance in our approach, and an emphasis on variance may keep small or moderate-effect QTL at intermediate frequency, at which variance is maximized (Lynch and Walsh 1998). Therefore, when selecting on CPM, we might expect a short-term sacrifice of genetic gain for long-term benefit. Indeed, we note a small deficit in selection response in the first two breeding cycles relative to FM selection, particularly for the trait under direct selection (Figure 5B). Selecting crosses using FM likely emphasizes the rapid increase in frequency of beneficial alleles at large and moderate-effect QTL, leading to fixation of unfavorable small-effect QTL due to drift or linkage (Figure 7B). Practically, the maintenance of genetic variance under CPM selection suggests that genetic gain may be sustained beyond 10 breeding cycles (Figure 5).

Changes in LD and haplotype frequencies are likely driving the movement towards zero of the genetic correlation in the breeding population. After the first breeding cycle, we observed a sharp trend towards, or persistence of, more negative genetic correlations (Figure 6B). This could be the product of negative covariance generated due to LD (Felsenstein 1965; Bulmer 1971; Falconer and Mackay 1996) and the simultaneous fixation of favorable haplotypes or QTL and maintenance of antagonistic haplotypes or QTL at intermediate frequency (Bennett and Swiger 1980; Falconer and Mackay 1996). Though genetic correlations became similar at later cycles (Figure 6B), the less rapid movement towards zero of genetic correlations when selecting crosses using CPM is curious. This pattern, more apparent with positive *r*_*G*(0)_, may be due to similar forces influencing the genetic variance. Selection on CPM combined with stronger selection intensity weighs the predicted genetic correlation and genetic variance of each trait (Equations 4 and 5). As above, this would value the maintenance of segregating QTL in the population, in agreement with our observations (Figure 7B). Co-segregating QTL for both traits would impact the genetic covariance to a greater degree than the genetic variances (Bohren *et al.* 1966; Villanueva and Kennedy 1990), leading to the observed differences in genetic correlation (Figure 6B). We would expect the linkage maintaining covariance in the short-term to be broken down by recombination, which may help explain why the genetic correlation moved more rapidly towards zero under the loose linkage genetic architecture (Figure 6B).

The general trends in genetic correlation over cycles can be explained by the genetic architecture. With pleiotropy, the movement towards – or maintenance of – a negative correlation is due to the fixation of favorable QTL and presence of antagonistic QTL. Absent pleiotropy, the correlation is due entirely to LD (Lande 1984; Lynch and Walsh 1998), which is degraded by recombination, eventually moving the genetic correlation towards zero (Villanueva and Kennedy 1990). Though our results confirm this under the loose linkage and tight linkage architecture (Figure 6B), the final genetic correlation is slightly negative, and more so with tight linkage. This is likely due to fixation of antagonistic QTL haplotypes, which, particularly when tightly linked, can effectively act as pleiotropic loci (Lande 1984) and are subject to the same competing forces mentioned earlier.

Our approach to cross selection for simultaneously improvement of multiple traits is one of several that have recently been proposed. Selection using CPM takes advantage of predictions of the genetic correlation between two traits (Equations 4 and 5). In a similar vein, Allier *et al.* (2019) proposed selecting crosses that were predicted to maximize the response for a trait under direct selection and produce the most favorable correlated response in parental contribution (treated as the indirect trait). Their method extends predictions of genetic correlations to three- and four-way crosses, a generalization beyond our described approach for biparental populations. Instead of predicting genetic correlations, a multi-trait index could be constructed and crosses could be selected on the basis of the superior progeny mean of that index (Yao *et al.* 2018). Though this simplifies the number of parameters to be predicted (a single genetic variance versus many pairwise correlations), it uses information differently from the CPM method. Finally, predictions of genetic correlations or correlated responses could enhance multi-objective cross optimization procedures (Akdemir *et al.* 2019).

The results of our simulation bode well for implementation in a breeding program. Notably, we observed that the advantage of selecting crosses on CPM was apparent even when the genetic correlation was negative (i.e. unfavorable). This condition is often encountered by breeders and would typically discourage the use of indirect selection (Bernardo 2010); however, we demonstrated that CPM cross selection can mitigate any negative response in the trait under indirect selection when the genetic correlation is negative.

### Application in a breeding program

To demonstrate its feasibility under more realistic conditions, we generated predictions of genetic correlations among populations in a barley breeding program. For two pairs of traits with moderately or strongly unfavorable correlations, we identified many crosses with favorable predicted correlations (Figure 4, Table 2), suggesting that specific crosses could be targeted to improve multiple traits simultaneously. This would rely on accurately discriminating among crosses, and we attempted to validate predictions of genetic correlations using empirical data of breeding populations.

Though we were only able to validate predictions for one pair of traits (Table 3), we observed that predictive ability seemed to be associated with the heritability of both traits, in agreement with the results from our first simulation. Of course, trait heritability may influence accuracy beyond unreliable marker effect estimates. With less heritable traits, the correlation among environmental effects is expected to have a greater influence on the phenotypic correlation (Lynch and Walsh 1998). It is not difficult to imagine how shared environment could influence the observed correlations. For instance, environmental stresses might stunt the growth of plants and promote earlier flowering, creating a positive correlation between these traits. Additionally, plants that flower later or are taller may avoid the soil-borne *F. graminearum* inoculum, potentially leading to artificial negative correlations between the traits. The results of our simulations and empirical experiment confirm that, as in any other implementation of genomewide selection, reliable phenotypic data is paramount.

The modest size of our TP (*n* = 175) likely constrained prediction accuracy, as suggested in our first simulation (Figure 2). Previous genomewide selection research, including those focused on barley, suggest a pattern of diminishing returns when predicting line means with ever-larger TPs (Lorenz *et al.* 2012; Sallam *et al.* 2015). A breeding program using a smaller TP, perhaps as a resource-saving measure, may be ill-equipped to utilize predictions of genetic correlations. The size of our TP was a function of the early stage of implementing genomewide selection in the breeding program, and we might expect that as more individuals are phenotyped and genotyped, the size of the training dataset will become more satisfactory.

Fortunately, the barrier for a breeder to incorporate predictions of genetic correlations is low. First, the data required for predictions (phenotypes, marker genotypes, and a genetic map) are commonly available in many breeding programs. Second, it is relatively inexpensive, in both time and computing power, to generate such predictions. Thus, it is possible that this procedure can be an additional tool for breeders to make decisions. We note, however, that validating predictions of genetic correlations requires phenotypic data on many large families, leading to population sizes that are generally unrealistic for a breeding program (Bernardo 2010). It is likely that these predictions, if implemented, will not be routinely validated, unlike genomewide predictions of genotypic means, and this lack of feedback may prove discouraging for breeders. Nevertheless, further work is necessary to demonstrate empirically the utility of selecting crosses informed by predictions of genetic correlations, with emphasis on the response to selection for two potentially unfavorably correlated traits.

## ACKNOWLEDGEMENTS

We thank Ed Schiefelbein, Guillermo Velasquez, and Karen Beaubien for technical support during population development, phenotypic data collection, and genotyping. Thanks go to Ruth Dill-Macky for suppling *F. graminearum* inoculum for St. Paul and to Madeline Smith and Joseph Wodarek for managing the FHB trial in Crookston, MN. We are grateful to Austin Case, Jo Heuschele, John Hill Price, Ian McNish, Becky Zhong, and Alexander Susko for assistance and encouragement. Resources from the Minnesota Supercomputing Institute were used to complete this project. This research was supported by the U.S. Wheat and Barley Scab Initiative, the Minnesota Department of Agriculture, Rahr Malting Company, the Brewers Association, the American Malting Barley Association, and USDA-NIFA Grant #2018-67011-28075.

