## Supplementary material for "Multi-Trait Improvement by Predicting Genetic Correlations in Breeding Crosses": FileS1

**File S1: Captions for Figures S1-S4 and Table S1**

**Multi-Trait Improvement by Predicting Genetic Correlations in Breeding Crosses**

Jeffrey L. Neyhart, Aaron J. Lorenz, and Kevin P. Smith

**Supplemental Figure Captions**

**Figure S1.** Comparison of predicted genetic parameters of 100 random simulated bi-parental recombinant inbred line (RIL) families using either the *PopVar* R package (Mohammadi *et al.* 2015) or a deterministic formula (Equations 1, 2, and 3 in main text). We compared the (**A**) cross mean and (**B**) genetic variance for two simulated quantitative traits, as well as (**C**) the genetic correlation between traits. The solid line denotes *y* = *x*.

**Figure S2.** Distributions of the realized base genetic correlation (*r_G(0)_*) in each iteration of our first breeding simulation (Simulation 1), plotted separately by number of quantitative trait loci (nQTL) and genetic architecture (Loose Linkage, Tight Linkage, or Pleiotropy) and colored by the target base genetic correlation (-0.5, 0, or 0.5). Realized *r_G(0)_* (*n* = 1800) were pooled over trait heritabilities and model for marker effects, but separated by training population sizes (*N_TP_*) of (**A**) 150, (**B**) 300, (**C**) 450, (**D**) 600.

**Figure S3.** Prediction accuracy of genetic correlations with increasing training population size, different heritabilities ($h^{2}$) of trait 1 ($h_{1}^{2}$) and trait 2 ($h_{2}^{2}$), genetic architectures (loose linkage, tight linkage, or pleiotropy), number of QTL (30 or 100), and genomewide prediction model [BayesCπ or RRBLUP (ridge regression best linear unbiased prediction)]. Results are separated by a starting genetic correlation (*r_G(0)_*) of (**A**) -0.5, (**B**) 0, and (**C**) 0.5.

**Figure S4.** Effect plot of the bias of the predicted genetic correlation, covariance, trait 1 genetic variance, and trait 2 genetic variance under different genetic architectures (red, loose linkage; blue, tight linkage; green, pleiotropy). Effects were calculated from a model that also accounted for trait 1 and trait 2 heritability, number of quantitative trait loci, training population size, and starting genetic correlation. Bias was calculated as $\left( \bar{x}-\bar{X} \right)/\bar{X}$, where $\bar{x}$ was the average predicted parameter across potential crosses (i.e. genetic variance, covariance, or correlation), and $\bar{X}$ was the average expectation of the parameter.

**Supplemental Table Captions**

**Table S1.** Prediction accuracy average (and 95% confidence interval) for the genetic correlation (CorG) between two traits, along with the mean and genetic variance (VarG) for each trait, in simulated breeding crosses. Accuracy of all parameters was measured in response to varying trait heritabilities (H2), number of quantitative trait loci (nQTL), training population (TP) size, the starting base genetic correlation (Base CorG), the genetic correlation architecture, and the model used to estimate genomewide marker effects. Results are summarized over 100 simulation replications.
