## Supplementary figures and images for "Multi-Trait Improvement by Predicting Genetic Correlations in Breeding Crosses"

### FigureS1

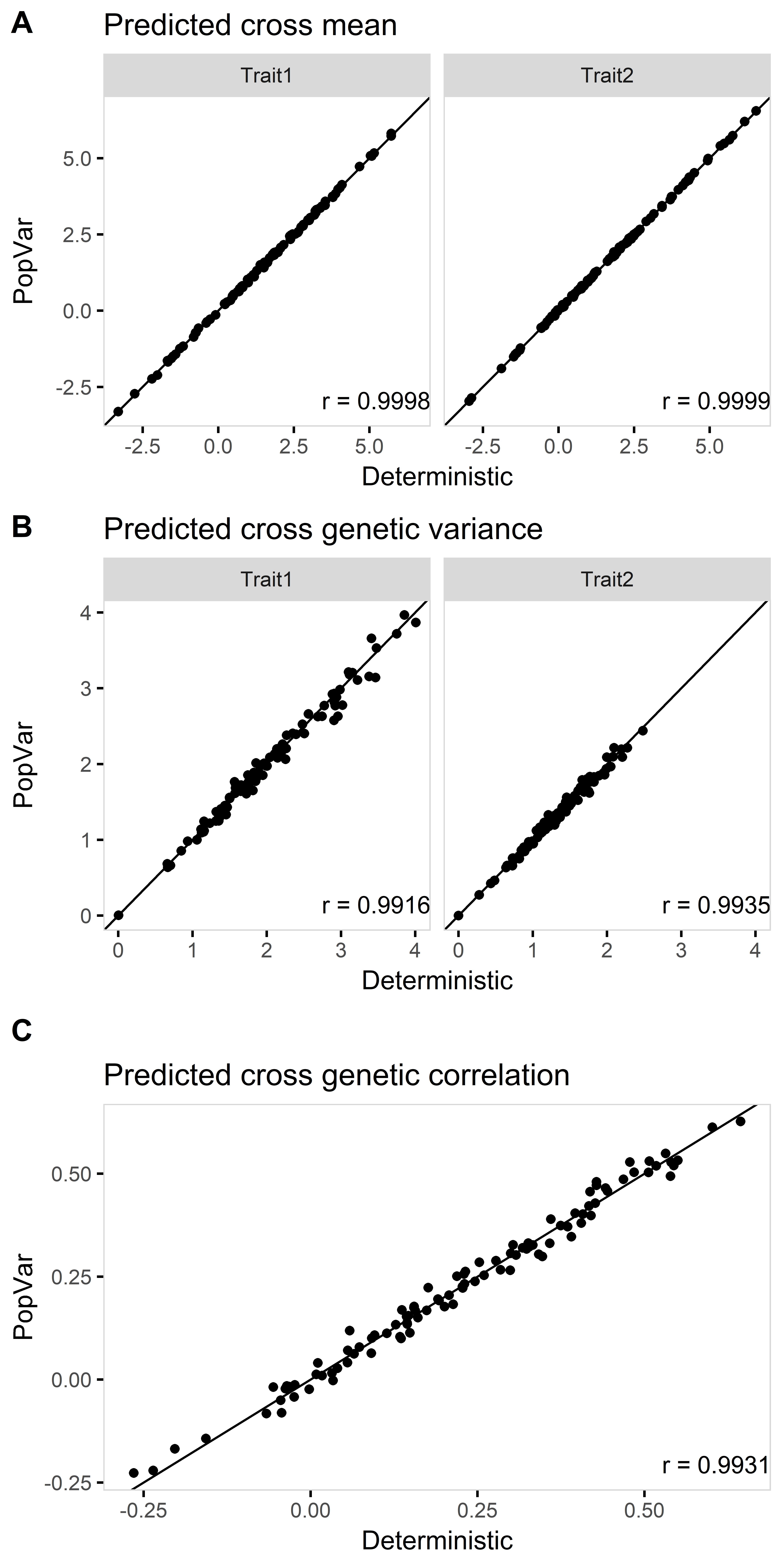

### FigureS2

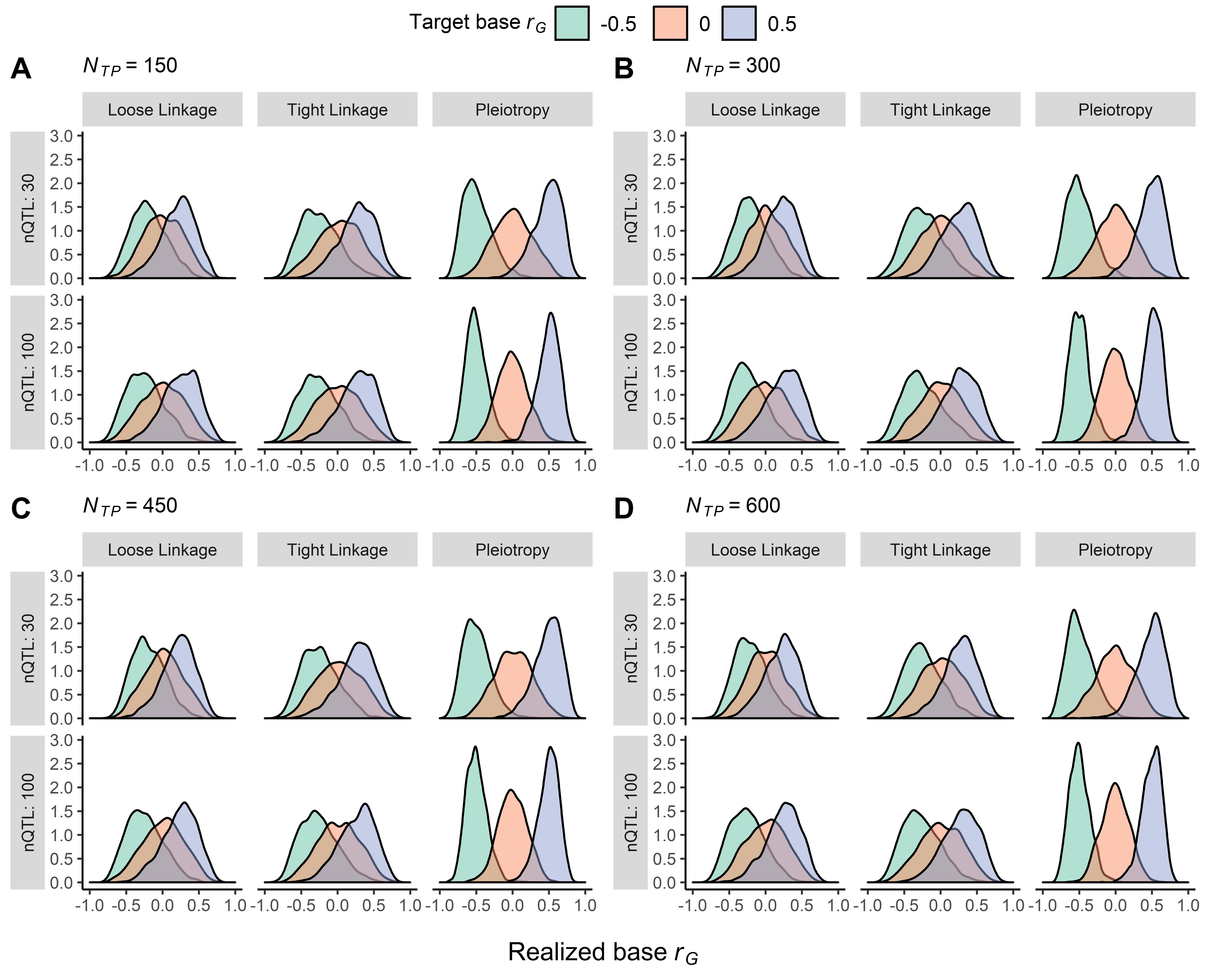

### FigureS3

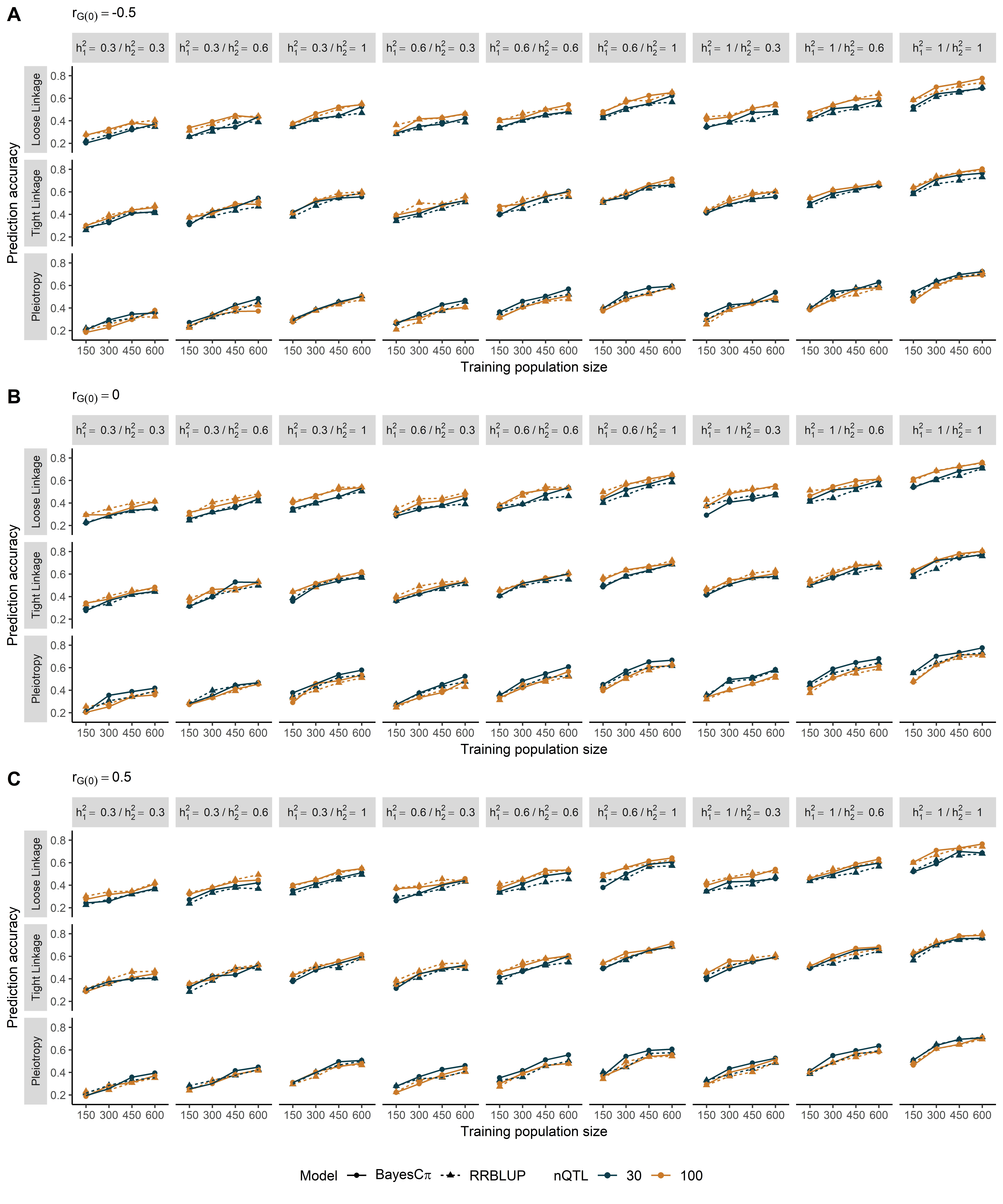

### FigureS4

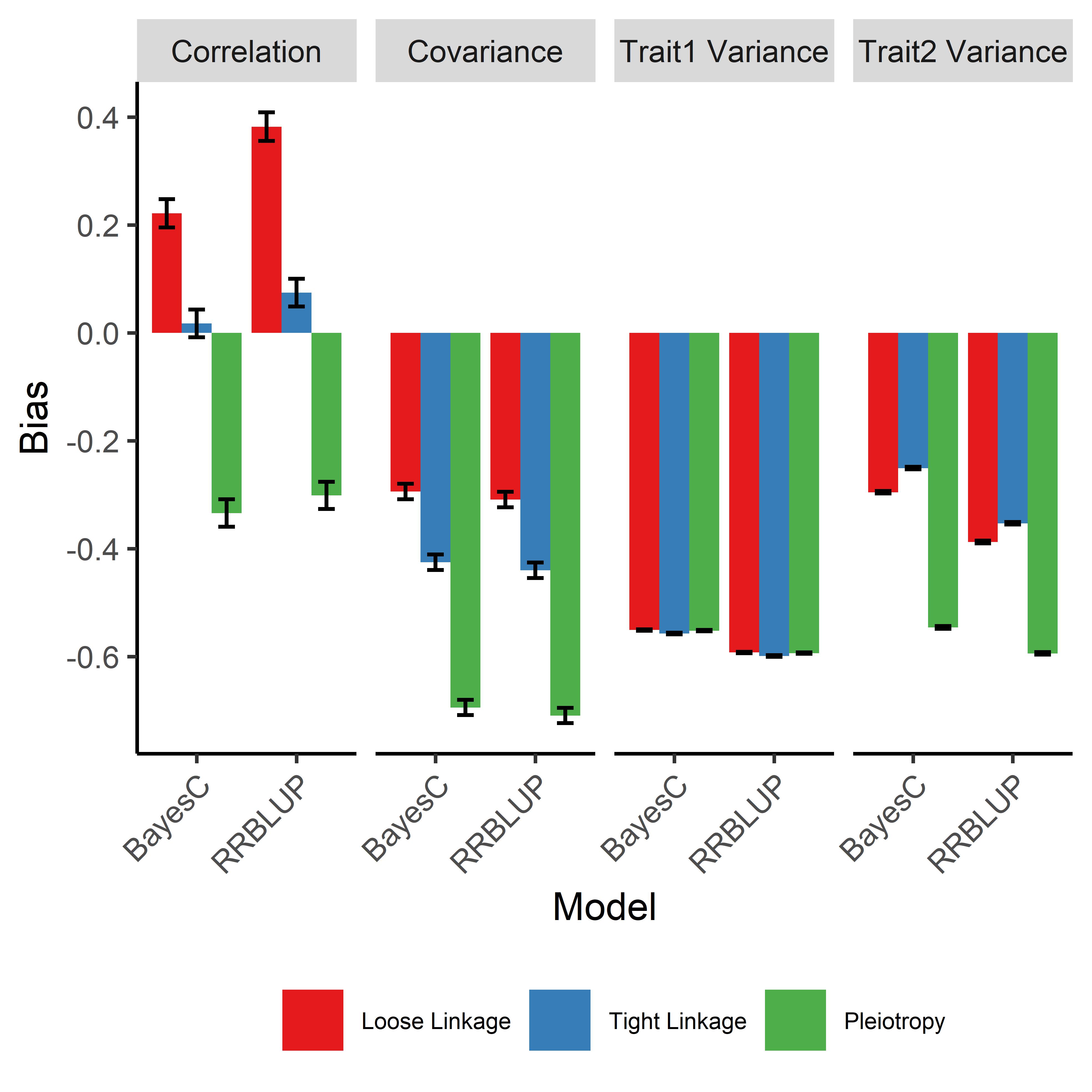
